# Community science for enigmatic ecosystems: Using eBird to assess avian biodiversity on glaciers and snowfields

**DOI:** 10.1101/2022.06.04.494825

**Authors:** William E. Brooks, Jordan Boersma, Neil Paprocki, Peter Wimberger, Scott Hotaling

## Abstract

**Aim:** To quantify avian biodiversity and habitat preference and describe behavior in an enigmatic, understudied ecosystem: mountain glaciers and snowfields.

**Location:** Mountains in the Pacific Northwest of western North America: British Columbia (CA), Washington and Oregon (USA).

**Taxon:** Birds observed within our study area and focal habitat.

**Methods:** We used community science data from eBird—an online database of bird observations from around the world—to estimate bird biodiversity and abundance in glacier and snowfield ecosystems as well as nearby, ice-adjacent habitats. We used field notes from eBird users and breeding codes to extend our data set to include insight into habitat usage and behavior. Finally, we compared our community-science approach to previous studies that used traditional survey methods.

**Results:** We identified considerable avian biodiversity in glacier and snowfield habitat (46 species) with four specialists that appeared to prefer glaciers and snowfields over nearby, ice-adjacent habitat. Combined with field notes by eBird users, our efforts increased the known global total of avian species associated with ice and snow habitats by 14%. When community science data was compared to traditional methods, we found similar species diversity but differences in abundance.

**Main conclusions:** Despite the imminent threat of glacier and snowfield melt due to climate change, species living in these habitats remain poorly studied, likely due to the remoteness and ruggedness of their terrain. Glaciers and snowfields hold notable bird diversity, however, with a specialized set of species appearing to preferentially forage in these habitats. Our results show that community science data can provide a valuable starting point for studying difficult to access areas, but traditional surveys are still useful for more rigorous quantification of avian biodiversity.

## Introduction

Contemporary climate change is driving global glacier recession reflected in mass loss and increased meltwater runoff (Bliss et al., 2014; Bolch et al., 2012; Moore et al., 2009). Recession of the mountain cryosphere–primarily glaciers and perennial snowfields–will negatively impact the considerable biodiversity these ecosystems host (Cauvy-Fraunié & Dangles, 2019; Hotaling, et al., 2021; Stibal et al., 2020). Glacier biodiversity is dominated by microbial communities thriving within, below, and downstream of glacial ice (Hotaling et al. 2017). A range of macroinvertebrates have also been described from glacier ecosystems, commonly acting as resource subsidies for larger taxa (e.g., birds, Hotaling et al., 2020). Finally, vertebrates also use glaciers for a host of activities including thermoregulation, food caching, and travel (Rosvold, 2016), but the extent of vertebrate links to glacier habitats remains poorly characterized. This knowledge gap is likely due to the rarity of vertebrates relative to smaller organisms paired with the remote, difficult-to-access terrain of glacier ecosystems, highlighting the need for alternative survey methods. Given the ongoing recession of the mountain cryosphere, more information about the biodiversity present in these habitats is urgently needed to quantify the impacts of glacier loss.

Among vertebrates, glaciers and perennial snowfields may represent particularly important habitats for birds. However, mountain birds are less studied than lowland species, despite widespread evidence for climate change impacting their reproduction, survival, and distributions (Scridel et al., 2018). While no systematic surveys of birds on glaciers and snowfields have been conducted, observations from literature reviews and a recent camera trap study have recorded just 22 bird species associated with glaciers worldwide (S. Hotaling, et al., personal communication, 2022; Goodman, 1971, Rosvold, 2016). Most birds appear to use cryospheric habitats for foraging on wind-blown arthropods, pollen, and seeds (Antor, 1995; Camfield et al., 2010; Crawford & Edwards, 1986; Hotaling et al., 2020; Shain et al., 2001). Several birds also nest near (Johnson, 1965; Rosvold, 2016) or on glaciers (Hardy & Hardy, 2008), likely due to food sources on the glacier surface and reduced predation risks.

Community science (also referred to as citizen science) is a rapidly growing source of data to fill gaps in ecological research (Kosmala et al., 2016; Silvertown, 2009). Community science uses volunteers to gather data, benefiting professional researchers through high-volume, low-cost data, and the community scientists through gained knowledge and exposure to research (Peter et al., 2021; Silvertown, 2009). Community science data have been successfully applied in biodiversity monitoring on large spatio-temporal scales (Callaghan et al., 2021; Chandler et al., 2017; Pocock et al., 2018), and to inform policy and decision-making (Conrad & Hilchey, 2011; de Sherbinin et al., 2021). The abundance of community science data presents opportunities to study rare species and habitats (Robinson et al., 2018), but these studies remain rare. For avian community science, eBird has emerged as a global leader; birdwatchers submit eBird “checklists”–i.e., lists of their field observations–containing bird abundances, qualitative observations, and sampling effort (Sullivan et al., 2009). These checklists can be aggregated to calculate trends and distributions of birds in locations or habitats of interest. In general, eBird-based estimates of abundances align well with traditional surveys (Callaghan et al., 2017; Callaghan & Gawlik, 2015).

In this study, we used eBird data to clarify avian biodiversity in an understudied, climate-change-threatened ecosystem: mountain glaciers and perennial snowfields. Through this objective, we present a framework for using a public, community science-derived database (eBird) to characterize links between a taxonomic group of interest (birds) and difficult-to-study habitats (glaciers and perennial snowfields in mountain ecosystems). Specifically, we compared avian communities and biodiversity in glaciers and snowfields and nearby, ice-adjacent habitats, and describe bird behavior in these habitats from observer notes. We show that eBird data can be used to rigorously quantify avian biodiversity and document habitat and behavior, highlighting the power of community science projects for studying uncommon and understudied ecosystems.

## Materials and Methods

### Data acquisition

We acquired eBird data (Sullivan et al., 2009) from alpine areas in southern British Columbia (BC), Washington (WA) and Oregon (OR) to document avian biodiversity on mountain glaciers and snowfields, as well as in nearby, ice-adjacent habitats. We chose this study region as it contains many mountain glaciers and is relatively highly populated, meaning it receives many eBird submissions compared to other alpine areas. We identified 129 focal alpine bird species (full list is included in Appendix 1) based primarily on two criteria: (1) they were detected in mountain bird surveys by Boyle and Martin (2015) or (2) have been reported in four frequently-used, high-altitude (2,149–3,157m elevation) eBird hotspots representative of our study region: Panorama Point (Mount Rainier National Park, WA, 46.804407, -121.721005), Blackcomb Mountain (BC, 50.092500, -122.887778), Mt. Hood (Hood River Co., OR, 45.372360, -121.672211), South Sister summit (Lane Co., OR, 44.102726, - 121.769070). eBird hotspots are public eBird locations created by eBird users that generally reflect popular birding locations. Species reported at each hotspot are aggregated and summarized on eBird.org. Additionally, based on known high-altitude migration in Baird’s sandpiper (*Calidris bairdii*; Shewey & Blount, 2017) and ice worm foraging by semipalmated plovers (*Charadrius semipalmatus;* Goodman, 1971), 12 common inland shorebird species were included to reflect the possible use of glaciers and snowfields during migration.

Collectively, we analyzed three separate data types reported by eBird users: observations, field notes, and breeding codes. Observation data are counts of each species reported by users, field notes are entered by users in a free text field for each observation, and breeding codes are optionally entered by users to document specific breeding behaviors. To acquire the data, we downloaded the “basic” eBird data set containing all eBird checklists submitted worldwide in June 2021. We filtered the data set in R version 4.1.0 using the auk_filter command of the auk package (RStudio Team, 2020; Strimas-Mackey et al., 2018) to only include checklists submitted in our focal region (WA, OR, and BC) that were marked as “complete.” Users mark their checklists as “complete” if they have reported all of the species they observed at a location. These complete checklists are ideal for measuring species diversity and abundance because they reduce overreporting of species considered more interesting to birdwatchers (e.g. rare species) and they can be used to infer the absence of species (those not reported). We then extracted complete checklists that included any of our 129 focal species.

### Spatial filtering

We spatially filtered our data set to bin checklists by those close to or including glaciers and snowfields versus nearby, ice-adjacent habitats. We considered both glaciers and snowfields together following Rosvold (2016). We included ice-adjacent habitat as a biodiversity comparison and to determine if any species had preference for glaciers and snowfields.

We used QGIS (QGIS development team, 2022) to create polygon layers defining our two habitat types. We used a permanent ice polygon layer to outline glaciers and snowfields (Natural Resources Canada et al., 2004), and added a 100m buffer around each polygon as it seemed likely that eBird users would observe birds from the edge of a glacier or snowfield. We created “ice-adjacent” habitat polygons with a 2-km buffer around all glaciers and snowfields which resulted in a layer roughly equal in area to the glacier and snowfield layer. We used the select by location tool to identify eBird checklists intersecting either of our habitat categories and extracted elevation data for each checklist from a 1-km resolution elevation raster layer using the point sampling tool (U.S. Geological Survey & Natural Resources Canada, 2007).

Next, we binned checklists into “glaciers and snowfield” or “ice-adjacent” habitat categories. We used the auk_zerofill command to add zeros where a species wasn’t observed, allowing each checklist to represent both presence and absence. We elected to remove checklists from northern British Columbia (latitude > 54.7° N) as glaciers transition to sea level there, thus possibly supporting a different bird community than the montane glaciers and snowfields that we were focused on. Since stringent filtering, particularly by sampling effort, generally improves accuracy in community science analyses (Steen et al., 2019; Van Eupen et al., 2021), we further filtered our zero-filled observation data to include only checklists that were stationary or with a travel distance under 1 km (85% reduction in total checklists). We chose a 1 km travel distance as it is half of the minimum width of most of our habitat polygons, making it less likely that users traveled out of the desired habitat.

### Calculating biodiversity metrics and comparing habitat types

To assess biodiversity in our focal habitats, we calculated species richness, or the total number of bird species observed, and Shannon-Weiner diversity index (H’). Before calculating Shannon-Wiener diversity index (H’ = -∑[(p_i_) × ln(p_i_)]), we removed 171 non-numeric observation (4% of all observations) which likely reflect community scientists marking species as present with an X, but not providing an abundance estimate. To identify the most common species, we calculated the observation frequency of each species as the proportion of all checklists where a given species is present.

We compared biodiversity and community composition in glaciers and snowfields to nearby, ice-adjacent habitat. After filtering, our data set consisted of 70 glacier and snowfield checklists and 887 ice-adjacent habitat checklists (Fig. 1). Since this 10:1 ratio ice-adjacent to glacier and snowfield checklists could bias our estimates of species richness, we resampled with replacement checklists for ice-adjacent habitats 10,000 times in sets of 70 and calculated species richness for each resampled set. Next, glacier and snowfield richness was compared to the 95% confidence interval (95th percentile) of the bootstrapped data from ice-adjacent habitat. We compared species diversity in glaciers and snowfields and ice-adjacent habitat for Shannon-Wiener indices with a Hutcheson t-test (Hutcheson, 1970), using an online calculator (https://www.dataanalytics.org.uk/comparing-diversity/).

**Figure 1.**
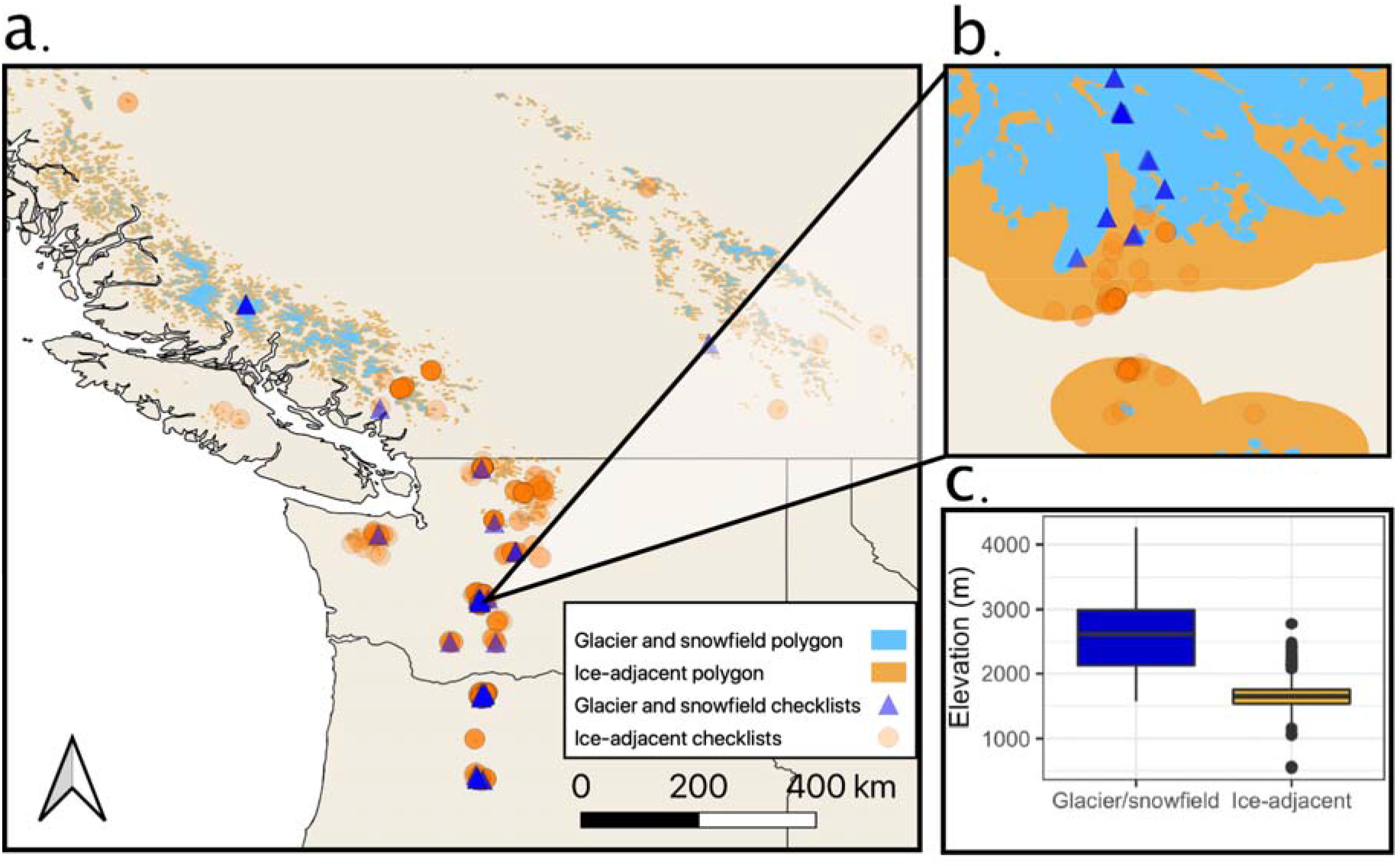
(a) Locations of eBird checklists on glaciers and snowfield (blue triangles) versus ice-adjacent habitats (orange circles). (b) Inset of the area around one of our focal hotspots—Panorama Point in Mount Rainier National Park, WA—showing how checklists were identified with habitat polygons. (c) Elevation of eBird checklists in our data set by habitat type.

To assess community similarity between glaciers and snowfields and ice-adjacent habitat, we compared species observation frequency in each habitat with linear regression. Normality of species frequencies was tested and corrected in ice-adjacent habitats with a cube root transformation. We also compared communities with Sorensen’s Coefficient (CC; Sorensen 1948) using the formula CC = 2 x Species in common / (Species count_1_ + Species count_2_). Finally, we examined habitat preference by subtracting the observation frequency of each species in nearby, ice-adjacent habitat from glaciers and snowfields; positive values represented species more common in glaciers and snowfields. We performed a series of Pearson’s Chi square tests to test for significant habitat associations.

### Using observer notes to understand habitat usage and behavior

We used observer field notes as an additional method for identifying glacier and snowfield habitat use by birds. eBird users report a wide array of field notes that represent a largely untapped source of descriptive data (Slager, 2020). We identified field notes containing habitat descriptions that included the keywords *glacier* and *snowfield* as well as variations of each (e.g. *snowfield* or *snow field*). We manually sorted field notes to remove false positives, thus only counting descriptions of birds in the habitat, not near it. We summed observations with notes for each bird species.

We assessed bird behavior on glaciers and snowfields using two previously defined data sets for glaciers and snowfields, checklists identified in our (1) spatial analysis and (2) field notes survey. We extracted descriptions of behavior using the same methods as habitat; we used keywords to identify relevant notes and manually reviewed them. We examined birds flying over glaciers and snowfields because they may be associated with the habitat despite not contacting snow or ice. We summed birds flying over (keyword: *flyover*) or using the flyover (F) breeding code. Similarly, since unseen birds heard by observers may be vocalizing from a glacier and snowfield, or nearby, ice-adjacent habitat we summed heard-only species in field notes (keywords: *heard only*). Finally, to examine how glaciers and snowfields function as a nutrient source, we identified foraging behavior (keywords: *feed, eat, forage*) and food choice in field notes (keywords: *spider, bug, insect, seed, pollen, worm*, and *ice worm*), and used the carrying food (CF) breeding code. We specifically sought descriptions of ice worm foraging, as this is a well-established but understudied resource for glacier-associated birds (Hotaling et al., 2020).

### Comparing community science data to traditional surveys

To compare diversity estimates generated by community science with traditional methods (e.g., transects) we performed two analyses. (1) We compared glacier- and snowfield-associated species identified in eBird field notes to those identified by Rosvold (2016) and a camera trap study of glacial wildlife (Hotaling et al. 2022). We chose these two studies as Rosvold (2016) was globally focused and Hotaling et al. (2022) was published later and focused on a glacier within our study area (Paradise Glacier, Mount Rainier, WA, USA). (2) We quantified the similarity of community composition and observation frequencies for our eBird data and traditional survey study—Boyle and Martin (2015)—for the ice-adjacent habitat. We were unable to do a similar analysis for snow/ice species because there are no traditional surveys of avian diversity for that habitat. We used Boyle and Martin (2015) as our comparison data set because it shared similar habitat, region, and total survey time [eBird “ice-adjacent”: ∼723 observation hours; Boyle and Martin (2015): 717 observation hours]. We compared the two data sets as described above using CC, observation frequencies, and linear regression.

## Results

Our initial data download included 1,984,111 eBird checklists. After spatial filtering, we identified 957 mountain checklists, comprising ∼845 observation hours of survey time. Mountain checklists were not evenly distributed in space or time: dates spanned 1999-2021 but were biased to summer months in the past 10 years. Geographically, most locations were in Washington (54%) and Oregon (40%) with few in British Columbia (6%; Fig. 1). We identified 70 checklists from glaciers and snowfields (∼122 observation hours) and 887 checklists from ice-adjacent habitats (∼723 observation hours). We extracted 867,989 species field notes from the total data set for text analysis, 52 of which contained keywords pertaining to habitat, behavior, or diet. We extracted just one observation with a breeding code relevant to our study.

### Avian biodiversity and community composition

Glacier and snowfield checklists contained 46 species (Fig. 2), and a Shannon-Weiner diversity index (H’) of 2.77. The most common species were common raven (*Corvus corax;* 47% of checklists), gray-crowned rosy-finch (*Leucosticte tephrocotis*; 46%), American pipit (*Anthus rubescens*, 29%), pine siskin (*Spinus pinus*, 11%), Clark’s nutcracker (*Nucifraga columbiana*, 10%), and dark-eyed junco (*Junco hyemalis*, 9%). All other species were reported in ≤ 6% of checklists. Overall, glacier and snowfield habitat exhibited lower biodiversity than nearby, ice-adjacent habitat. We identified 106 species in the ice-adjacent data set. However, after correcting for sample size, ice-adjacent habitats were still more biodiverse but the difference was much smaller at ∼68 ice-adjacent species (bootstrapped 95% CI: 61-76; Fig. S1). Glaciers and snowfields also had lower Shannon-Weiner diversity (H’ = 2.77) than ice-adjacent habitats (H’ = 3.66; Hutcheson’s t-test: t = 14.69, p < 0.001).

**Figure 2.**
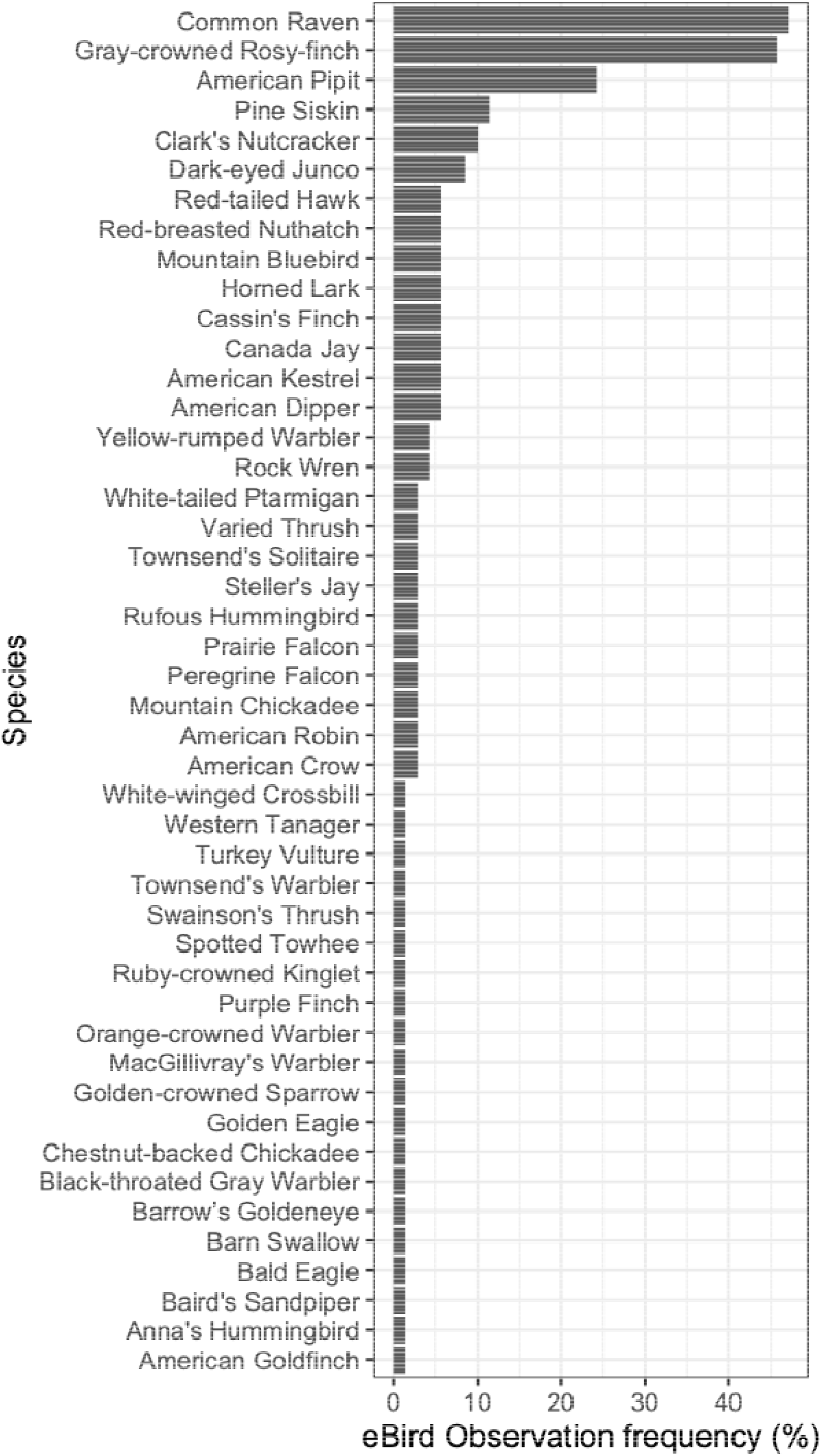
Observation frequency for all birds included in our data set for glacier and snowfield habitat.

Community composition was similar in glaciers and snowfields versus ice-adjacent habitats: roughly 59% of species were present in both habitats (CC = 0.59) and observation frequencies were significantly correlated (R^2^ = 0.17, F_1,127_ = 26.59, p < 0.001). Four species were significantly more common in glaciers and snowfields than ice-adjacent habitats (Fig. 3): gray-crowned rosy-finches (Pearson’s Chi Square test, *X*^*2*^ = 161, p < 0.001), American pipits (*X*^*2*^ = 10.6, p = 0.001), rock wren (*X*^*2*^ = 5.61, p = 0.018), and American kestrel (*X*^*2*^ = 5.09, p = 0.024). All were evenly distributed across our study region except rock wren, which was primarily reported in Oregon.

**Figure 3.**
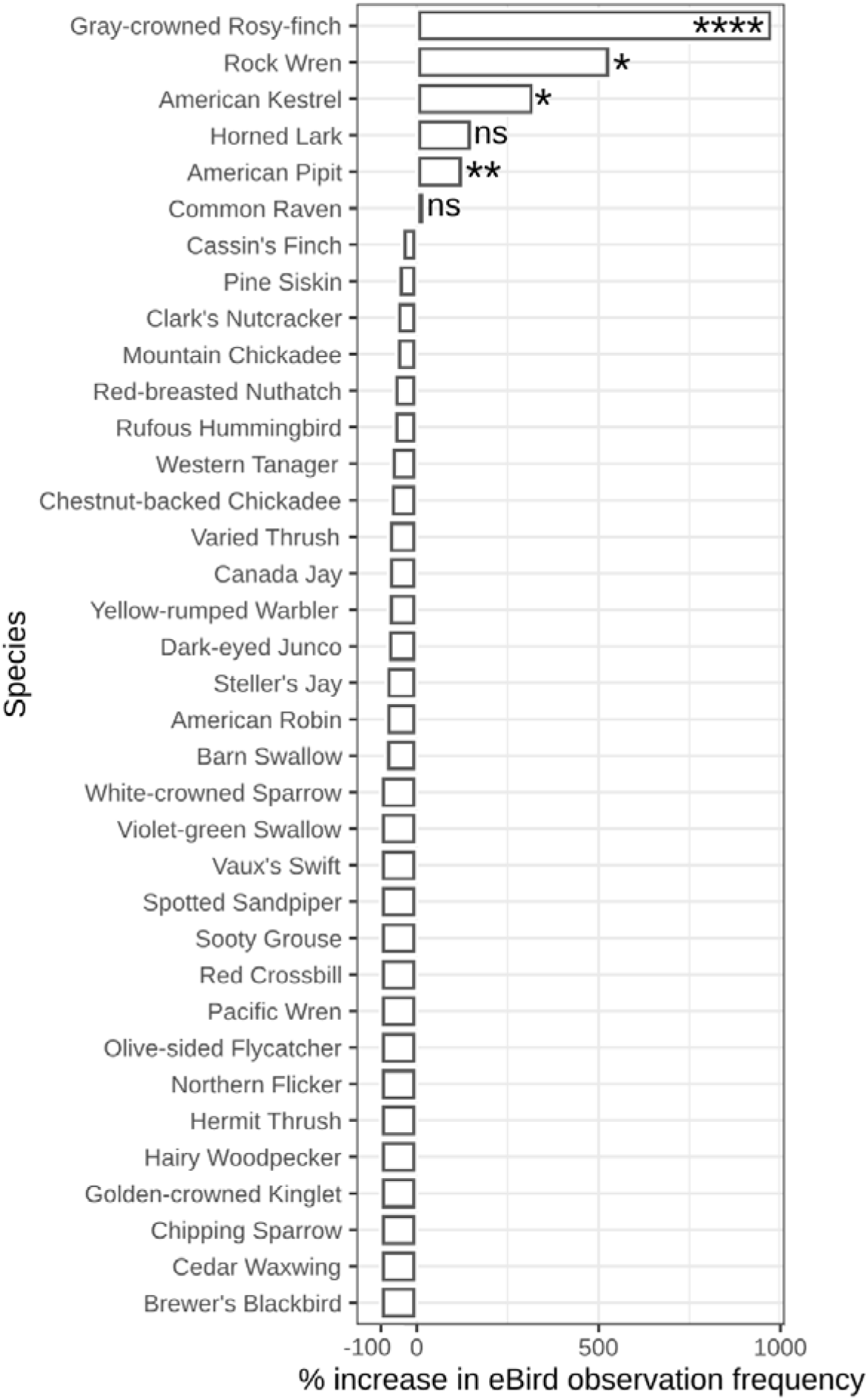
The difference in eBird observation frequency (%) for each species in glaciers and snowfield habitat relative to ice-adjacent habitat. Statistical significance from chi-square tests indicate species that are more common in glaciers and snowfields at p < 0.0001 (****), p < 0.001 (***), p < 0.01 (**), and p < 0.05 (*). Non-significant values are indicated with “ns.” Only observation frequency differences ≥ 0.03 are shown.

### Observer-reported habitat and behavior

We identified field notes describing 12 species occupying mountain glaciers and snowfields (Table 1). Travel by flying over glaciers and snowfields was noted for 13 species, and ground travel was only observed for white-tailed ptarmigan (*Lagopus leucura*). Common ravens were observed sliding down snowfields on their breasts. Feeding on glaciers and snowfields was noted for five species with arthropods, conifer seeds, and ice worms noted as food sources (Table 1). Baird’s sandpipers were the only species observed feeding on ice worms. Observations of predators feeding in glaciers and snowfields were rare. A golden eagle (*Aquila chrysaetos*) was observed feeding on something on a snowfield, and a prairie falcon (*Falco mexicanus*) was observed chasing American pipits near a glacier. American pipits (3 obs.) and a yellow-rumped warbler (*Setophaga coronata*; 1 obs.) carrying food were the only observations marked with a breeding code. eBird users described hearing, but not seeing, five species: Clark’s nutcracker (2 obs.), Cassin’s finch (*Haemorhous cassinii*, 3 obs.), dark-eyed junco (1 obs.), pine grosbeak (*Pinicola enucleator*, 1 obs.), and yellow-rumped warbler (1 obs.).

**Table 1.**
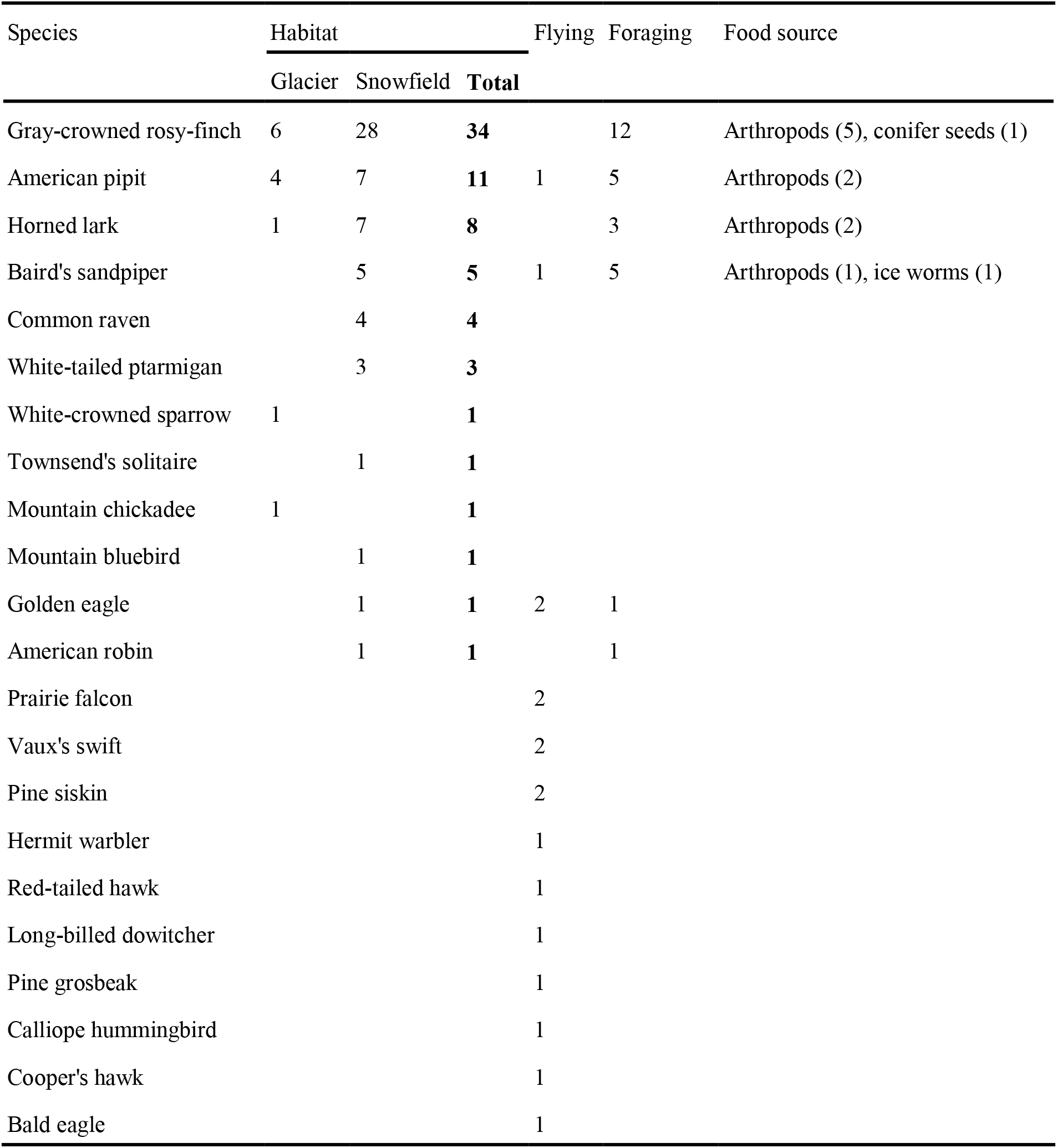
Habitat and behavior descriptions extracted from eBird field notes. Birds flying over glaciers and snowfields were counted separately from birds standing in either habitat. The number of observations for each food source is shown in parentheses.

### Community-science versus traditional field surveys of avian biodiversity

Field notes documented species occupying glaciers and snowfields that matched previous observations: 7 of 19 species compiled in a global review (Rosvold, 2016) and all 7 species documented by camera trapping at Mount Rainier, Washington (Hotaling et al. 2022). We identified four new glacier- or snowfield-associated species using community-science data: Baird’s sandpiper, white-crowned sparrow (*Zonotrichia leucophrys*), mountain chickadee (*Poecile gambeli*), and Townsend’s solitaire (*Myadestes townsendi*). Our community science data set was also comparable to a traditional transect-based survey approach for ice-adjacent habitat. Our data set shared 80% of species (CC = 0.81) with Boyle and Martin (2015). Moreover, eBird data outperformed traditional survey methods by capturing 11 more species. Observation frequency of each species was moderately correlated in both data sources (Fig. 4; Linear regression: R^2^ = 0.35, F_1,130_ = 71.02, p < 0.001).

**Figure 4.**
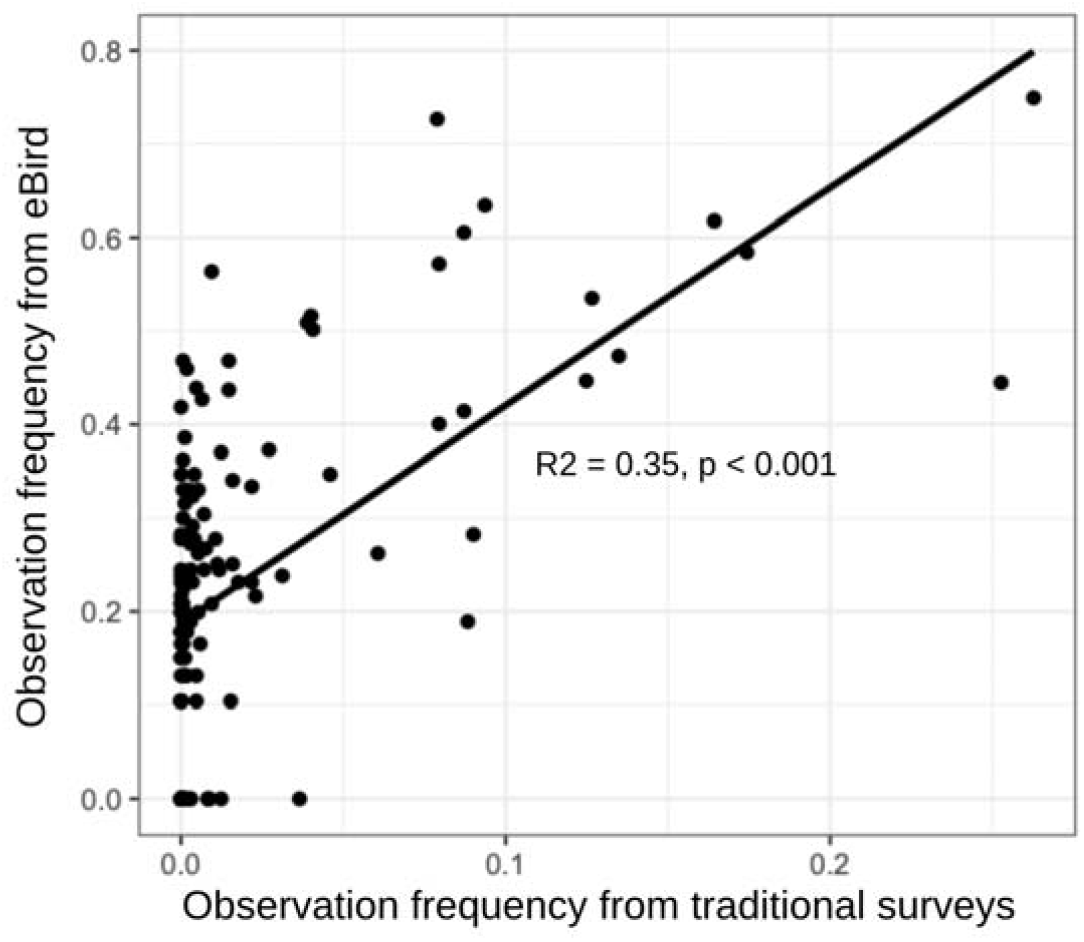
Species observation frequencies for community science data (eBird) versus traditional surveys (Boyle and Martin 2015). Each point represents a single species.

## Discussion

Mountain glaciers and snowfields support unique, specialized biodiversity but are critically threatened by climate change worldwide (Hotaling et al., 2020; Stibal et al., 2020). To date, vertebrate biodiversity on glaciers and snowfields has been understudied (Rosvold, 2016), likely due to difficulty of access. In this study, we used community science to characterize avian diversity on glaciers and snowfields, highlighting the power of this type of data for improving understanding of hard-to-study ecosystems. Through community-science data, we identified 46 bird species associated with glaciers and snowfields including four species that had not been associated with these habitats previously. Given the limited geographic scope of our study, we expect that new cryosphere-associated species remain to be discovered via community science approaches.

### Specialized avian biodiversity on mountain glaciers and snowfields

Our results demonstrate strong preference in gray-crowned rosy-finches and American pipits for glacier and snowfield habitats which aligns with prior studies (e.g., Rosvold 2016), including observations of gray-crowned rosy-finches nesting near glaciers (Johnson, 1965). Other species more common in these habitats were more surprising, notably American kestrels and rock wrens. American kestrels have also been observed via camera trapping (Hotaling et al., 2022), and are known to occupy a broad range of open habitats, including alpine tundra. Rock wrens have not been documented using glaciers or snowfields but are known to be high-elevation tolerant (Benedict et al., 2021). Collectively, our results suggest that a small, overlooked community of ice specializing birds may exist, likely with region-specific taxa (e.g., white-winged diuca finches nesting in crevasses of Andean glaciers, Hardy & Hardy, 2008).

### Perennial snow and ice as a resource subsidy for mountain birds

Glaciers and snowfields capture drifting material like arthropods and seeds, and in some regions, hold *in situ* nutrient sources like ice worms (Hotaling et al., 2017), thereby providing easily obtainable food sources for alpine birds (Antor, 1995; Crawford & Edwards, 1986; Edwards & Banko, 1976; Mann et al., 1980; Rosvold, 2016). Glaciers and snowfields appear to serve as a nutrient supplement during two energetically-demanding life-cycle periods: breeding and migration (Hotaling et al. 2020). Most of the species observed foraging by eBird users are alpine breeders, so glaciers and snowfields likely hold important nutrient subsidies early in the breeding season (June) when snow cover reaches its peak (Hotaling et al., 2020). One species, Baird’s sandpiper, was only recorded as a migrant in July and August. We captured the first observation of Baird’s sandpipers feeding on ice worms, adding to six bird species previously recorded (Hotaling et al., 2020). The use of glaciers and snowfields by higher trophic level species remains poorly understood (Rosvold, 2016), but we identified some evidence of hunting by birds of prey. Hunting habitat choice may explain apparent preference by American kestrels for glaciers and snowfields as smaller birds may be easy to target against a white background.

### North American avian biodiversity imperiled by glacier melt

Across species and space, a general rule of the biodiversity impacts of glacier recession on ice-linked species emerges: as glaciers decline, the abundance of habitat generalists increase and specialists decline (Cauvy-Fraunié & Dangles, 2019). For instance, American Pipits showed the second strongest preference for glaciers and snowfields in our study, often using the ice and snow for foraging. In a well-studied population in Wyoming, up to 18% of foraging trips were made to snowfields during the breeding season (Hendricks, 1987; Hendricks & Verbeek, 2020). However, American Pipits also seem to prefer foraging on south-facing slopes, which are typically more snow free (Norvell & Creighton, 1990). It is possible that glacier melt will negatively impact American Pipit populations by reducing food availability during breeding, but their use of diverse feeding habitat suggests they may be able to supplement their diet in adjacent habitats.

Among North American species, gray-crowned rosy-finches, however, may be most at risk from glacier melt. In addition to a strong preference for foraging on glaciers and snowfields, gray-crowned rosy-finch are the highest-nesting songbird in North America (MacDougall-Shackleton et al., 2020).

Species distribution modeling indicates that they are extremely vulnerable to projected climate change in the next half century due to their narrow, high-altitude breeding niche (Conrad, 2015). It seems likely that loss of foraging habitat will be a major factor, particularly during breeding, contributing to population declines. Our evidence and previous observations (Hotaling et al., 2020; Rosvold, 2016) suggest gray-crowned rosy-finches frequently feed on arthropods and ice worms in glaciers and snowfields, and that invertebrate feeding increases during breeding (MacDougall-Shackleton et al., 2020). Whether gray-crowned rosy-finches can adjust their diet to compensate for changing food resources is unclear.

### Leveraging community science to study enigmatic ecosystems

While potential for sampling bias is ever present in eBird data (Boyle & Martin, 2015; Sullivan et al., 2009), we were able to extract a large sample size to characterize avian communities in difficult-to-study mountain habitats. We found species that matched previous observations from glaciers and snowfields, and our data set was comparable in terms of community composition and species richness to traditional approaches. However, to generate comparable data sets and mitigate sampling bias, it is important to normalize data, perhaps by using resampling methods like the bootstrapping approach we applied. In a similar study focused on a rare species—tricolored blackbird (*Agelaius tricolor*)— undersampling of absence data was used to compensate for sampling bias and improve species distribution model performance (Robinson et al., 2018). In our view, community science data represent a highly valuable resource for studying uncommon species and habitats as long as appropriate measures are taken to address sampling bias.

We used eBird field notes to extend our data set to include behavioral insight. eBird users more frequently described behavior in field notes rather than breeding codes. Considering the vast eBird framework, a more automated approach to processing field notes could provide equally powerful insights for behavior as the existing observation-focused framework. Such automation could be achieved by applying natural language processing (NLP; Hirschberg and Manning 2015), which uses computational methods to interpret written language. NLP has been successfully applied in other fields (e.g., health outcomes from medical records; Velupillai et al. 2018), and pre-trained models have emerged recently, which simplify the application of NLP methods (Qiu et al., 2020).

The most difficult challenge to using community science data for our research goals was overcoming spatial uncertainty in eBird data. eBird checklists with a shorter travel distance produce more precise location data, and thus higher spatial certainty (Steen et al., 2019). Therefore, we faced a common tradeoff between spatial resolution and data abundance for community science data when rare species or understudied habitats are considered: more stringent filtering by travel distance meant a smaller data set (Van Eupen et al., 2021). Additionally, user error when submitting eBird checklist locations and/or inclusion of species heard or seen in other areas may falsely inflate records. At present, eBird data is a valuable starting point for studying difficult to access areas or assessing large-scale (e.g., global) patterns, but traditional surveys are still required when rigorous quantification is needed.

Beyond birds, similar community science analyses of rare and under-studied ecosystems can likely be conducted for other taxa. Many platforms aggregate data from non-avian taxa, but some have structural features that limit data quality and value. For example, iNaturalist is a popular online platform that accepts reports of any organism (or abiotic evidence of organisms, e.g., tracks). The broad scope of iNaturalist allows a remarkable range of creative studies, like testing for increased use of urban habitat by mammals during the COVID-19 pandemic lockdown (Vardi et al., 2021). However, projects using iNaturalist data are limited by presence-only data. Because eBird checklists have the option to be marked as complete, absence data and abundance estimates can be inferred, thereby leading to conclusions that are more robust and resistant to sampling bias (Robinson et al., 2018). Additionally, iNaturalist uses a community reviewing system, with observations deemed “research grade” when enough users have confirmed the identification. The community reviewing system includes rewards for number of reviews, creating a potential incentive for maximizing quantity over quality when reviewing identifications. For instance, research grade observations have been shown to not have higher taxonomic accuracy (Hochmair et al., 2020), suggesting a community review system is not sufficient for data validation. Adoption of a system similar to eBird’s—where rare species are captured by an automatic filter and reviewed by pre-established experts—could improve this issue.

## Conclusion

Community science data clearly provide a valuable resource for quantifying biodiversity and engaging non-professional scientists in research. In this study, we described a framework for using eBird to assess avian biodiversity in understudied mountain cryosphere habitats that are imperiled by climate change. While supporting lower avian diversity than nearby, ice-adjacent habitats, glaciers and snowfields appear to provide key foraging grounds for many birds, including a few specialized species. As climate change proceeds, we expect that a loss of foraging grounds may negatively impact species like the gray-crowned rosy-finch. As community science platforms grow and mature, we expect their research value will continue to improve and eventually allow for robust temporal assessments of biodiversity change as year-over-year records accumulate. Collectively, our study highlights the power of community science data for modern ecological research, including the additive power that user reported field notes can add to observational data. We hope the professional research community continues to embrace community science data as we seek to realize their full potential for monitoring global biodiversity amidst an array of anthropogenic threats.

## Data availability

Data is available for download on eBird.org. All code used for this study is publicly available on GitHub (https://github.com/willbrooks0/eBird-glacier.git).

## Acknowledgements

We thank eBird developers, users, and reviewers for their contributions to the eBird project, and those who publish publicly available coding help.

## Biosketch

William Brooks received his BS in biology from the University of Puget Sound. As part of his undergraduate thesis he used eBird data to measure sparrow range expansion due to anthropogenic habitat change. In the future, he will examine bird responses to deforestation in tropical systems.

## Author contributions

W.B. and P.W. conceived of the study. W.B., J.B., N.P., P.W., and S.H. gave input on study design and data analysis. W.B. collected and analyzed the data. W.B. wrote the initial manuscript draft and all authors made considerable contributions to its development. All authors approved its submission. Authors declare no conflict of interest.

## Appendices

### Appendix 1 The 129 focal species included in this study

American Coot, American Crow, American Dipper, American Goldfinch, American Kestrel, American Pipit, American Robin, American Three-toed Woodpecker, American Tree Sparrow, Anna’s Hummingbird, Baird’s Sandpiper, Bald Eagle, Band-tailed Pigeon, Barn Swallow, Barrow’s Goldeneye, Black Swift, Black-bellied Plover, Black-billed Magpie, Black-capped Chickadee, Black-throated Gray Warbler, Bohemian Waxwing, Boreal Chickadee, Brewer’s Blackbird, Brown Creeper, Brown-headed Cowbird, California Scrub-Jay, Calliope Hummingbird, Canada Goose, Canada Jay, Cassin’s Finch, Cassin’s Vireo, Cedar Waxwing, Chestnut-backed Chickadee, Chipping Sparrow, Clark’s Nutcracker, Common Raven, Common Redpoll, Common Yellowthroat, Cooper’s Hawk, Dark-eyed Junco, Downy Woodpecker, Dusky Grouse, European Starling, Evening Grosbeak, Fox Sparrow, Golden Eagle, Golden-crowned Kinglet, Golden-crowned Sparrow, Gray-crowned Rosy-finch, Greater Yellowlegs, Hairy Woodpecker, Hammond’s Flycatcher, Hermit Thrush, Hermit Warbler, Horned Lark, House Finch, House Sparrow, Killdeer, Least Sandpiper, Lesser Yellowlegs, Lincoln’s Sparrow, Long-billed Dowitcher, MacGillivray’s Warbler, Mallard, Merlin, Mountain Bluebird, Mountain Chickadee, Mourning Dove, Nashville Warbler, Northern Flicker, Northern Goshawk, Northern Harrier, Northern Pygmy-Owl, Olive-sided Flycatcher, Orange-crowned Warbler, Pacific Wren, Pacific-slope Flycatcher, Pectoral Sandpiper, Peregrine Falcon, Pileated Woodpecker, Pine Grosbeak, Pine Siskin, Prairie Falcon, Purple Finch, Red Crossbill, Red-breasted Nuthatch, Red-breasted Sapsucker, Red-naped Sapsucker, Red-tailed Hawk, Rock Pigeon, Rock Wren, Rough-legged Hawk, Ruby-crowned Kinglet, Ruffed Grouse, Rufous Hummingbird, Savannah Sparrow, Semipalmated Plover, Semipalmated Sandpiper, Sharp-shinned Hawk, Snowy Owl, Solitary Sandpiper, Song Sparrow, Sooty Grouse, Spotted Sandpiper, Spotted Towhee, Spruce Grouse, Steller’s Jay, Swainson’s Thrush, Swamp Sparrow, Townsend’s Solitaire, Townsend’s Warbler, Tree Swallow, Turkey Vulture, Varied Thrush, Vaux’s Swift, Vesper Sparrow, Violet-green Swallow, Warbling Vireo, Western Bluebird, Western Meadowlark, Western Sandpiper, Western Tanager, White-crowned Sparrow, White-tailed Ptarmigan, White-winged Crossbill, Willow Ptarmigan, Wilson’s Warbler, Yellow Warbler, and Yellow-rumped Warbler.

