## Supporting Information for "Community science for enigmatic ecosystems: Using eBird to assess avian biodiversity on glaciers and snowfields"

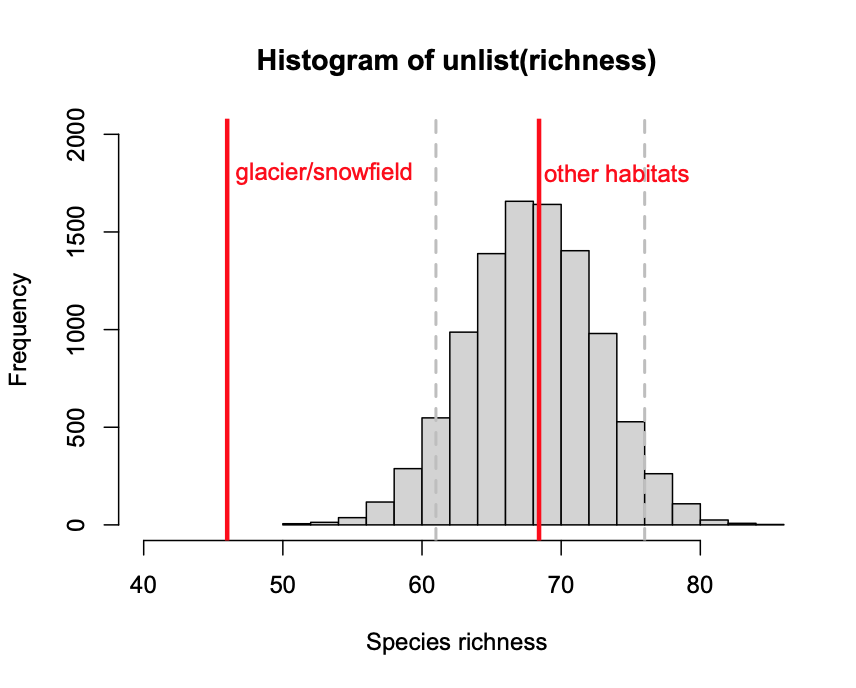

**Figure S1**. Species richness of glaciers and snowfields, and bootstrapped species richness of ice-adjacent habitat. The red lines indicate species richness for each habitat type. The dotted lines indicate 95% confidence intervals for bootstrapped data. Species richness is significantly lower in glaciers and snowfields versus ice-adjacent habitat.
